# Influenza Viruses in Mice: Deep Sequencing Analysis of Serial Passage and Effects of Sialic Acid Structural Variation

**DOI:** 10.1101/683516

**Authors:** Brian R. Wasik, Ian E.H. Voorhees, Karen N. Barnard, Brynn K. Lawrence, Wendy S. Weichert, Grace Hood, Aitor Nogales, Luis Martínez-Sobrido, Edward C. Holmes, Colin R. Parrish

**Author notes:** Center for Animal Health Research, INIA-CISA, 28130 Valdeolmos, Madrid, Spain.

## Abstract

Influenza A viruses have regularly jumped to new hosts to cause epidemics or pandemics, an evolutionary process that involves variation in the viral traits necessary to overcome host barriers and facilitate transmission. Mice are not a natural host for influenza virus, but are frequently used as models in studies of pathogenesis, often after multiple passages to achieve higher viral titers that result in clinical disease such as weight loss or death. Here we examine the processes of influenza A virus infection and evolution in mice by comparing deep sequence variation of a human H1N1 pandemic virus, a seasonal H3N2 virus, and a H3N2 canine influenza virus during experimental passage. We also compared replication and sequence variation in wild-type mice expressing N-glycolylneuraminic acid (Neu5Gc) with that seen in mice expressing only N-acetylneuraminic acid (Neu5Ac). Viruses derived from plasmids were propagated in MDCK cells and then passaged in mice up to four times. Full genome deep sequencing of the plasmids, cultured viruses, and viruses from mice at various passages revealed only small numbers of mutational changes. The H3N2 canine influenza virus showed increases in frequency of sporadic mutations in the PB2, PA, and NA segments. The H1N1 pandemic virus grew well in mice, and while it exhibited the maintenance of some minority mutations, there was no clear adaptive evolution. The H3N2 seasonal virus did not establish in the mice. Finally, there were no clear sequence differences associated with the presence or absence of Neu5Gc.

**SIGNIFICANCE:** Mice are commonly used as a model to study the growth and virulence of influenza A viruses in mammals, but are not a natural host and have distinct sialic acid receptor profiles compared to humans. Using experimental infections with different subtypes of influenza A virus derived from different hosts we found that evolution of influenza A virus in mice did not necessarily proceed through the linear accumulation of host-adaptive mutations, that there was variation in the patterns of mutations detected in each repetition, and the mutation dynamics depended on the virus examined. In addition, variation in the viral receptor, sialic acid, did not effect influenza evolution in this model. Overall this shows that mice provide a useful animal model for influenza, but that host passage evolution will vary depending on the virus being tested.

## INTRODUCTION

Many animal viruses naturally infect and spread among a range of host species, and can often be experimentally inoculated into alternative hosts to cause infections and disease, and where onward transmission may result in epidemics. Because the spread of a virus in a new host may involve adaptation to that host, with ongoing selection as the virus replicates in the host cells, tissues, and populations, or responds to host immunity (1–3), determining the underlying processes of viral evolution is central to understanding and controlling the spread of viruses in humans and other animal populations.

Here we combine the infection of influenza A virus (IAV) in mice with complete genome deep sequencing to examine the underlying processes and dynamics of viral growth in a new host. Most IAVs are naturally maintained as intestinal infections of various bird species in fresh and salt water environments (4). Occasionally, IAV will spill-over to infect mammalian hosts, and more rarely will go on to cause epidemics and pandemics. Natural outbreaks by IAV in mammals have been observed in humans, swine, mink, seals, horses, cats and dogs (5, 6). We have the greatest knowledge of outbreaks that occurred during the past 100 years, which in humans include the H1N1, H2N2, H3N2 human seasonal viruses that were first recognized in 1918, 1957, and 1968, respectively, as well as of the second human H1N1 pandemic strain that spread worldwide in 2009, replacing the circulating seasonal H1N1 clade (7, 8). The emergence of IAV in new hosts involves the natural selection of mutations involved in host adaptation, which may alter binding to the sialic acid (Sia) receptor by the Hemagglutinin (HA), cleavage of the Sia by the Neuraminidase (NA), host specific nuclear transport and replication processes that involve the polymerase subunits, and evasion of the host immune responses (reviewed by 2, 9–11). Some of these processes and mutations that impact mammalian transmission and disease have also been identified during experimental host passages, in which IAVs are passaged in new hosts such as ferrets (12–14).

Mice are not a natural host for IAVs, but have long been used as an animal model to study the replication, pathogenesis, and immune responses of many different viruses from avian and mammalian hosts (15–18). While some IAV strains appear to infect and replicate to high levels in the lungs or other respiratory tissues of mice, many show relatively limited replication, most do not spread naturally from mouse to mouse, and many cause little disease unless they are adapted by serial passage (19–21). The sequences of mouse-adapted IAVs often contain mutations in various genomic segments, including HA, NA, NS, as well as the polymerase gene segments (PB2, PB1, PA) (reviewed by 22). The considerable number of mouse adaptation studies of IAV vary in repetition, reproducibility, and specifics of the methodologies and experimental variables, all of which could impact virus evolution. In addition, analyses have frequently relied on laboratory tissue culture isolation and the measurement of population consensus mutations or polymorphisms that rose to high levels, often assuming that these were mouse-adaptive mutations fixed by positive selection. However, aside from a direct fitness advantage, some of the mutations observed in mice might have attained higher frequencies due to founder effects associated with population bottlenecks, and/or by hitchhiking with beneficial mutations. The acquisition of increased fitness in a new host may also involve complex epistatic mutations, and can depend on the specific sequences of the genomes in which they arise (genetic contingency) (23–25). Together, these issues indicate that endpoint analysis of genome variation in passaged populations may not distinguish functionally adaptive mutations from other non-adaptive variation (26). Understanding the details of how mutations arise in viruses during passage in mice would therefore provide a better understanding of the intricate evolutionary processes involved, and facilitate a comparison to viral emergence seen in other natural hosts.

Viral receptors are often involved in host adaptation, and influenza viruses use Sia-terminated glycans on cell glycoproteins or glycolipids as primary receptors of infection while also interacting with and evading or removing Sia in the mucus of the respiratory tract (27, 28). Sia are a family of glycans that include N-acetylneuraminic acid (Neu5Ac) as well as other modified forms, and are primarily connected to the underlying glycan through α2,3 or α2,6 linkages (α2,8 linkages are primarily found in polysialic acids) (29). Modifications of Sia may include the hydroxylation of the 5-acetyl group to glycolyl by the cytidine monophosphate-N-acetylneuraminic acid hydroxylase (CMAH), creating N-glycolylneuraminic acid (Neu5Gc) (30). Humans and ferrets lack a functional CMAH so that Neu5Gc is not displayed in cells of those hosts, while CMAH is active in other natural influenza hosts such as swine and horses, and Neu5Gc is also displayed at high levels in many tissues of mice (31, 32). Other chemical modifications of Sia that have the potential to impact virus-Sia interactions are also abundant and diverse across vertebrate species, and include the addition of acetyl groups on the 4, 7, 8 and/or 9 positions, as well as potential lactyl or sulfate modifications of the glycerol side chain (33–35). IAV emergence in new hosts frequently involves adaptation to the specific linkage of the Sia in the respiratory tissues of that host (36). In one common process, the HAs of avian IAVs bind with higher affinity to α2,3-linked Sia receptors, which are more abundant in the avian gastrointestinal tract (37). In contrast, human IAV HAs preferentially bind the α2,6-linked Sias that predominate the human upper respiratory tract (URT), and adaptation to humans involves adaptive mutation of the HA receptor binding site (36, 38, 39).

Mice differ in a number of properties compared to the natural hosts of IAV that include birds, humans, dogs, swine, horses, seals and mink. Their laboratory utility in pathogenesis and immunology is often in contrast to ferrets and guinea pigs that are commonly used as experimental model of virus contact and airborne transmission (40, 41). Mice also often differ in the forms of the natural Sia receptor, as they display lower proportions of α2,6 linked Sia in their URT relative to humans and some other hosts, and display both Neu5Ac and Neu5Gc (42, 43). The α2,3- and α2,6-linked Sias and other differences can result in selection in both the HA1 and HA2 domains (44). The HA and NA of some influenza strains may distinguish between Neu5Ac and Neu5Gc, generally through lower binding of the HA and lower NA activity on Neu5Gc compared to Neu5Ac (45, 46). Mice that lack Neu5Gc have been generated by knocking-out the CMAH gene which was then crossbred into a C57BL/6 background (31), allowing the specific biology of that variation to be assayed, as well as their effects on the Sia specificity of pathogenic bacteria (47, 48).

Here we use three different IAVs - human viruses A/California/04/2009 H1N1 and A/Wyoming/3/2003 H3N2, and canine A/Canine/IL/11613/2015 H3N2 - to define, in detail, the sequence variation that arises during infection and passage in mice. Each virus was derived from a genetically homogeneous starting point in reverse genetics plasmids, propagated only a limited number of passages in cells to prepare an infectious virus stock, and then passed in wild-type C57BL/6 mice and in *CMAH*^−/−^ mice that lack Neu5Gc. We used full genome deep sequencing to define the experimental variation of the viruses, and the role of potential Neu5Gc receptors in that process.

## RESULTS

### The IAVs tested were strain-specific for replication and phenotypic pathology in mice

The A/California/04/2009 H1N1 CA04 (H1N1p), A/Wyoming/3/2003 H3N2 WY03 (H3N2hu) and A/Canine/IL/11613/2015 H3N2 IL15 (H3N2ca) influenza viruses were each derived from plasmid clones (**Fig. 1A**). The H1N1p and H3N2ca were recovered and passaged in MDCK cells, while the H3N2hu were recovered and grown in MDCK-SIAT cells expressing higher levels of α2,6-linked Sia. Analysis of wild-type C57BL/6 mice revealed that Neu5Gc comprised between 45% and 60% of total the Sia present in the trachea and lungs (**Table 1**). The *CMAH*^−/−^ mice contained no detectable levels of Neu5Gc within the same tissue samples (**Table 1**; **Fig. 1B**). Viruses were inoculated into wild-type C57BL/6 mice or *CMAH*^−/−^ mice, then transferred through serial lung-to-lung passages with a fixed volume of 50µl (**Fig. 1C**). The inoculations with H1N1p and H3N2ca were transferred through four groups of C57BL/6 and *CMAH*^−/−^ mice in the first series of passages, and through three groups in the second series of passages (**Fig. 2**).

**Figure 1.**
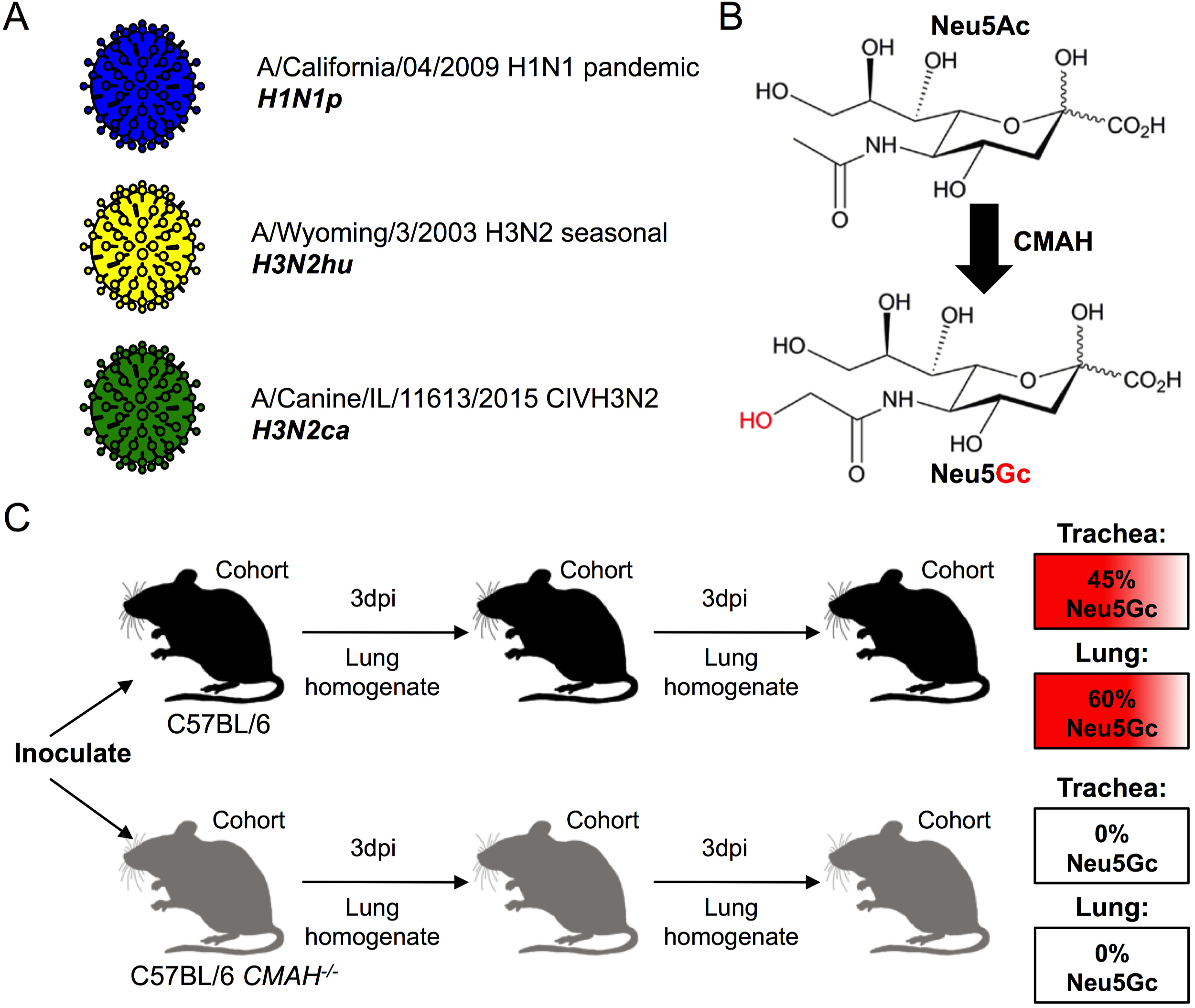
Outline of the experimental design. (A) Influenza virus stocks were generated from reverse genetic clones: H1N1p (blue), H3N2hu (yellow), H3N2ca (green). (B) The enzyme CMAH catalyzes the enzymatic conversion of Neu5Ac to Neu5Gc. Inactivation of the *cmah* gene in a C57BL/6 mouse background generates a Neu5Ac-only mouse. (C) Experimental passages of viruses were performed by nasal inoculation of culture-derived virus or lung homogenates into control C57BL/6 (black) or *CMAH*^−/−^ (grey) mice cohorts, 3-day incubation, and harvest of lung homogenates. C57BL/6 mice display Neu5Gc, as a proportion of total Sia, at 45% in trachea and 60% in the lungs. *CMAH*^−/−^ mice contain no Neu5Gc in their trachea or lungs.

**Figure 2.**
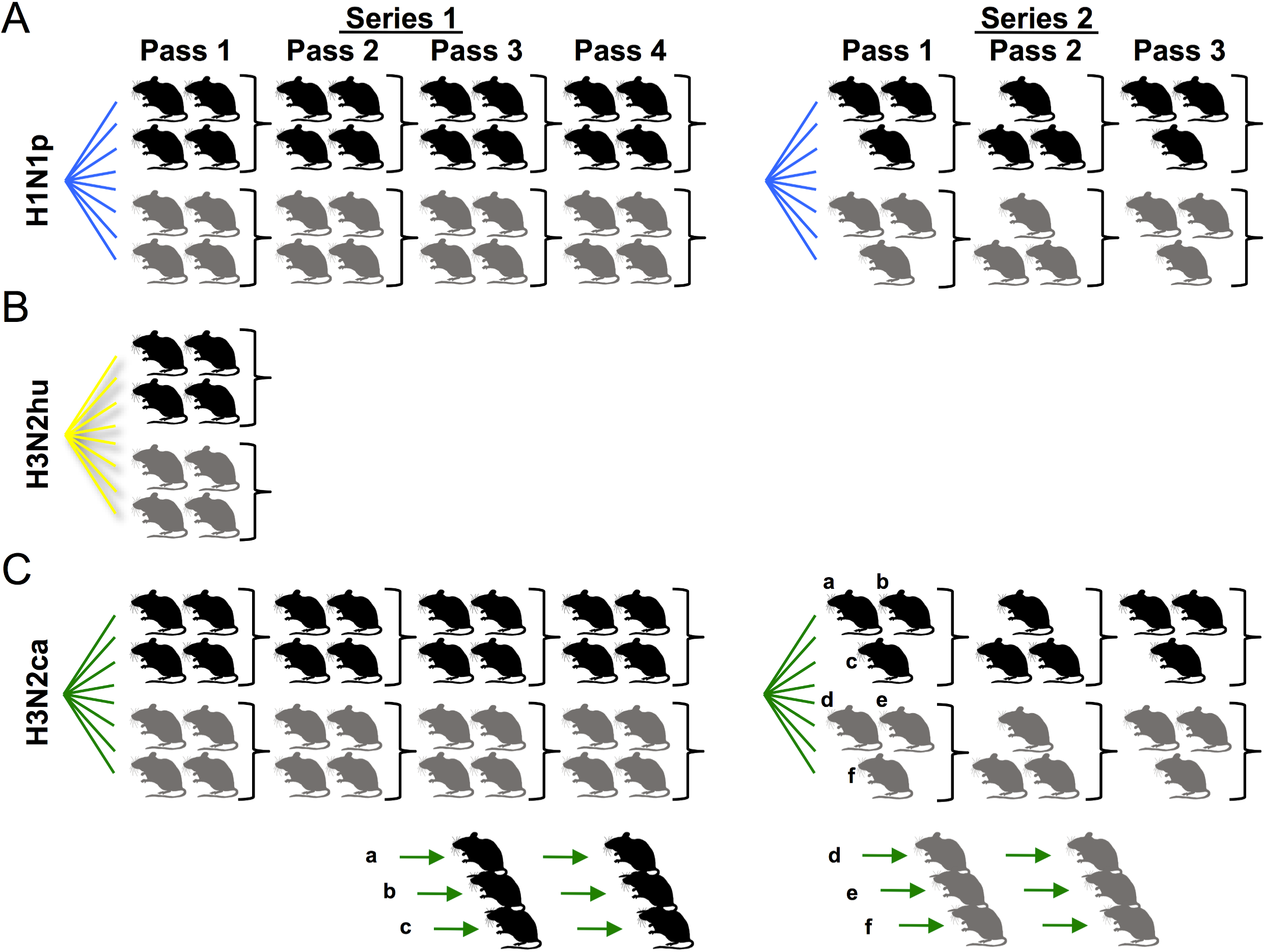
Diagram of experimental mouse passages performed in this study with H1N1p (A), H3N2hu (B), H3N2ca, or H3N2ca in mouse-to-mouse lineages (C). The experiment was performed in two different iterations: Series 1 proceeded for four passages among cohorts of four mice (two male, two female), while Series 2 proceeded for three passages among cohorts of three mice (an alternating 2:1 sex ratio). H3N2ca mice lung homogenates from passage 1 of Series 2 were also used to initiate a series of mouse-to-mouse lineages in C57BL/6 (a, b, c) or *CMAH*^−/−^ (d, e, f) individual mice.

**Table 1.**

|  | C57BL/6 Mice |  | *CMAH*<sup>-/-</sup> Mice |  |
| --- | --- | --- | --- | --- |
|  | Neu5Ac | Neu5Gc | Neu5Ac | Neu5Gc |
| **Trachea** | 50% | 45% | 92% | n/d |
| **Lung** | 40% | 60% | 96% | n/d |

The relative proportion of Neu5Ac and Neu5Gc (of total Sia) identified in mouse respiratory tissues, as determined by HPLC analysis. n/d = not detected.

Mice were initially inoculated with a fixed number of TCID_50_ units of virus as measured in MDCK or MDCK-SIAT cells, while genomic vRNA quantitation revealed that doses varied between 10^6^ and 10^8^ genome copies per 50 µL inoculum. The H1N1p virus replicated to give robust genome copy numbers maintained in lungs per passage (generally 2-3 log_10_ units more than the initial dose), and mice displayed moderate to severe signs of disease including weight loss, observed lethargy, and the presence of lesions on the lungs (**Fig. 3A**). In contrast mice inoculated with seasonal human H3N2hu virus (**Fig. 3B**), even at 10^8^ genome copies, showed few genome copies in their lungs upon the first passage, and exhibited no observable signs of infection including weight loss. No virus was recovered after transfer of materials of the lungs of the first passage mice to a second series of mice (data not shown). The H3N2ca virus (**Fig. 3C**) replicated well in the lungs of mice during the experiments, although with 2 or 3 log_10_ lower titers present in lungs after the initial passage. The H3N2ca virus infected mice showed occasional behavioral signs of infection during the 3-day incubation (lethargy), but exhibited no measured weight loss and no notable anatomical abnormality of the respiratory tissues.

**Figure 3.**
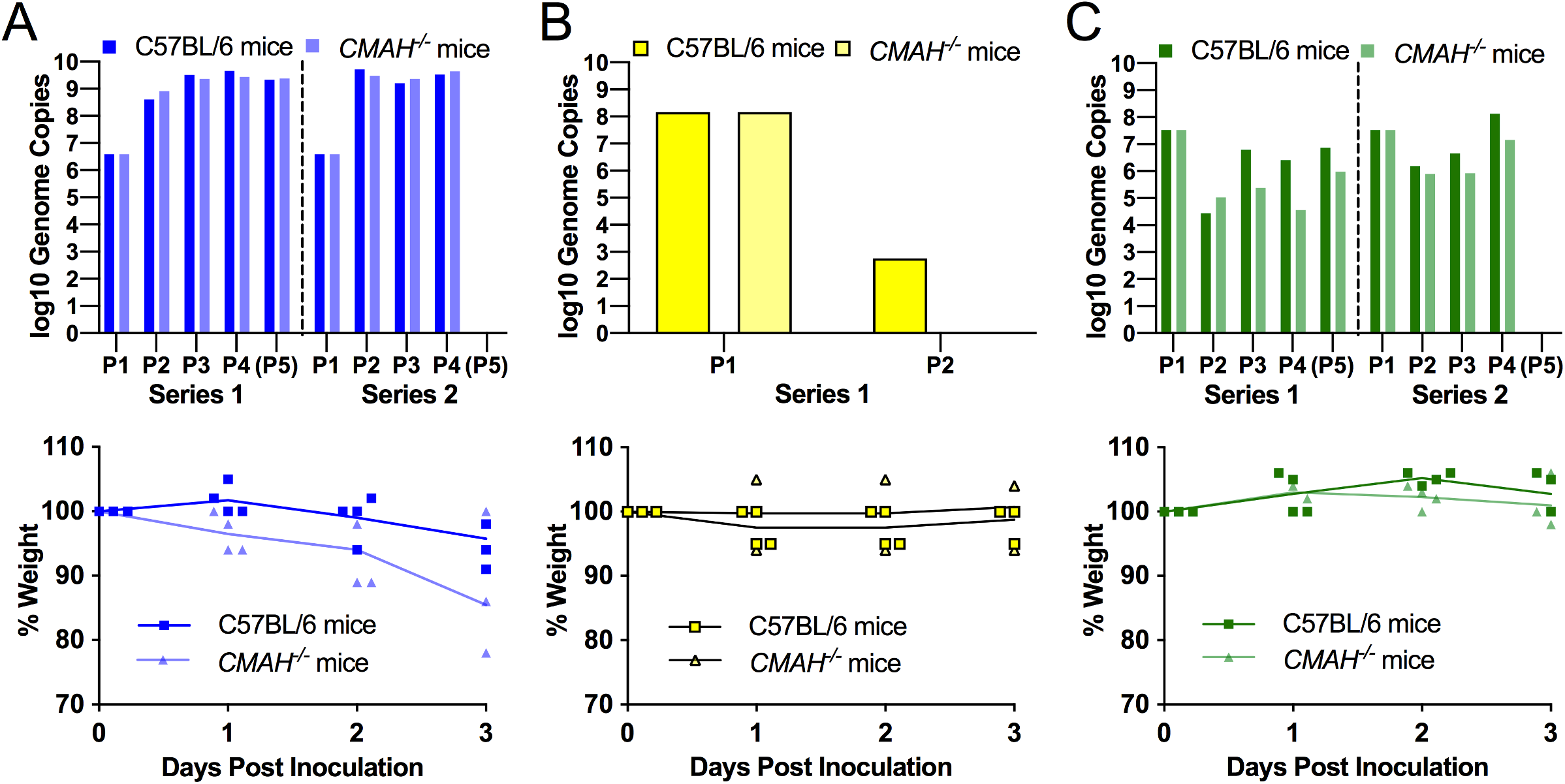
General influenza dynamics in mouse cohorts inoculated with H1N1p (A), H3N2hu (B), or H3N2ca (C). Top, quantitation of influenza genome copies (per RT-qPCR of M segment) for each virus was measured for stock inoculum and pooled lung homogenate inoculum (50µl) to measure genomic bottleneck size at each passage. H3N2hu-inoculated mice lacked measurable genome copies in their lungs at first passage. Bottom, mouse weights were recorded during course of infection. Figures are from the first passage of Series 1 as a representative example. All virus-specific weight-loss phenotypes persisted during course of each experimental passage and between repeat passage series. Only mice inoculated with H1N1p showed weight loss during course of infection, with no significant variation between experimental groups.

We examined the respiratory tissues of C57BL/6 and *CMAH*^−/−^ mice to determine whether alterations in Neu5Gc expression would impact the qualitative display of Sia receptor linkages. Histochemistry with the lectins MAH and SNA to detect either α2,3- or α2,6-linked Sias showed no obvious differences in the staining of those tissues (**Fig. 4A**). Immunohistochemistry of the mouse lungs confirmed the presence of IAV antigen (NP) in H1N1p- and H3N2ca-inoculated mice, but not in H3N2hu-inoculated mice (**Fig. 4B**).

**Figure 4.**
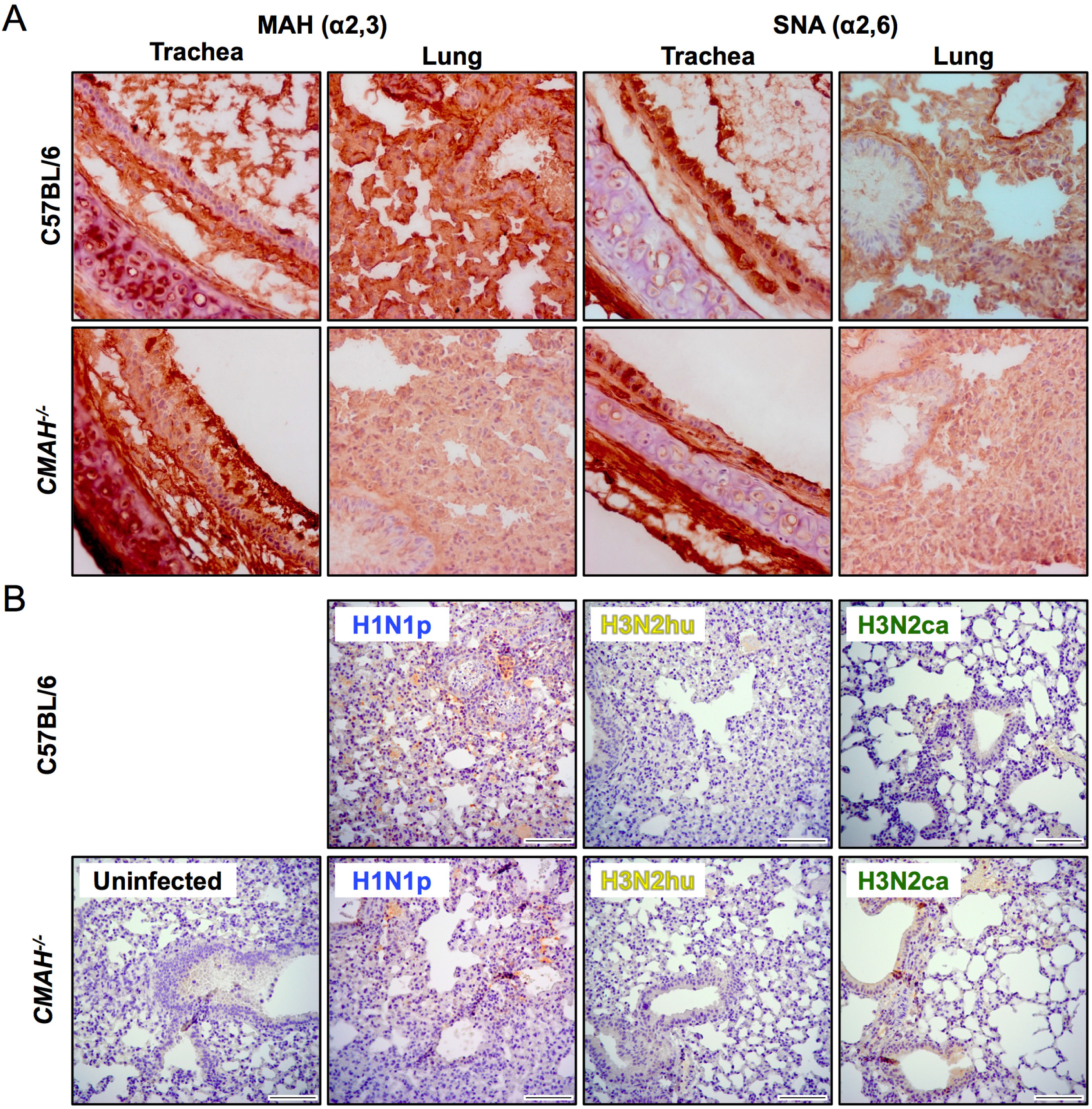
(A) Expression of the α2,3 - and α2,6-linked Sia in the trachea and lungs of wild-type C57BL/6 mice that express ∼45-60% Neu5c, or *CMAH*^−/−^ mice, which lack Neu5Gc. (B) Examples showing the viral infection in the lungs of experimentally passaged mice by immunohistochemistry for IAV antigen (NP), stained in red. Scale bar = 100 µm.

Viral RNA (vRNA) were amplified by IAV whole-genome RT-PCR from pooled and individual experimental lungs (as outlined in **Fig. 2**), and the cDNA used to generate libraries for Illumina sequencing. RT-qPCR of the M segment confirmed that genome copies were generally >100,000 RNA copies per reaction, allowing more confident analysis of low percentage variants (49). The median copy number per reaction of H1N1p samples was 4.15 × 10^8^ (±2.15 × 10^8^), while samples of H3N2ca showed greater variation in amounts, although with a medium number of copies of 1.99 × 10^5^ (±2.15 × 10^6^) per reaction, although even the least abundant sample was in excess of 4000 RNA copies (**Fig. 5A**). Amplification and read coverage varied among segments despite robust quality vRNA in all reactions, and the PB2/PB1/PA segments frequently showing several fold lower mean coverage per base compared to other segments for the same virus strain (**Fig. 5B**). This may be driven by inconsistent coverage across the length of PB2/PB1/PA segments (**Fig. 5C**). We set stringency cut-offs for single nucleotide variant (SNV) calling at 1.0% with minimum coverage of 500 reads.

**Figure 5.**
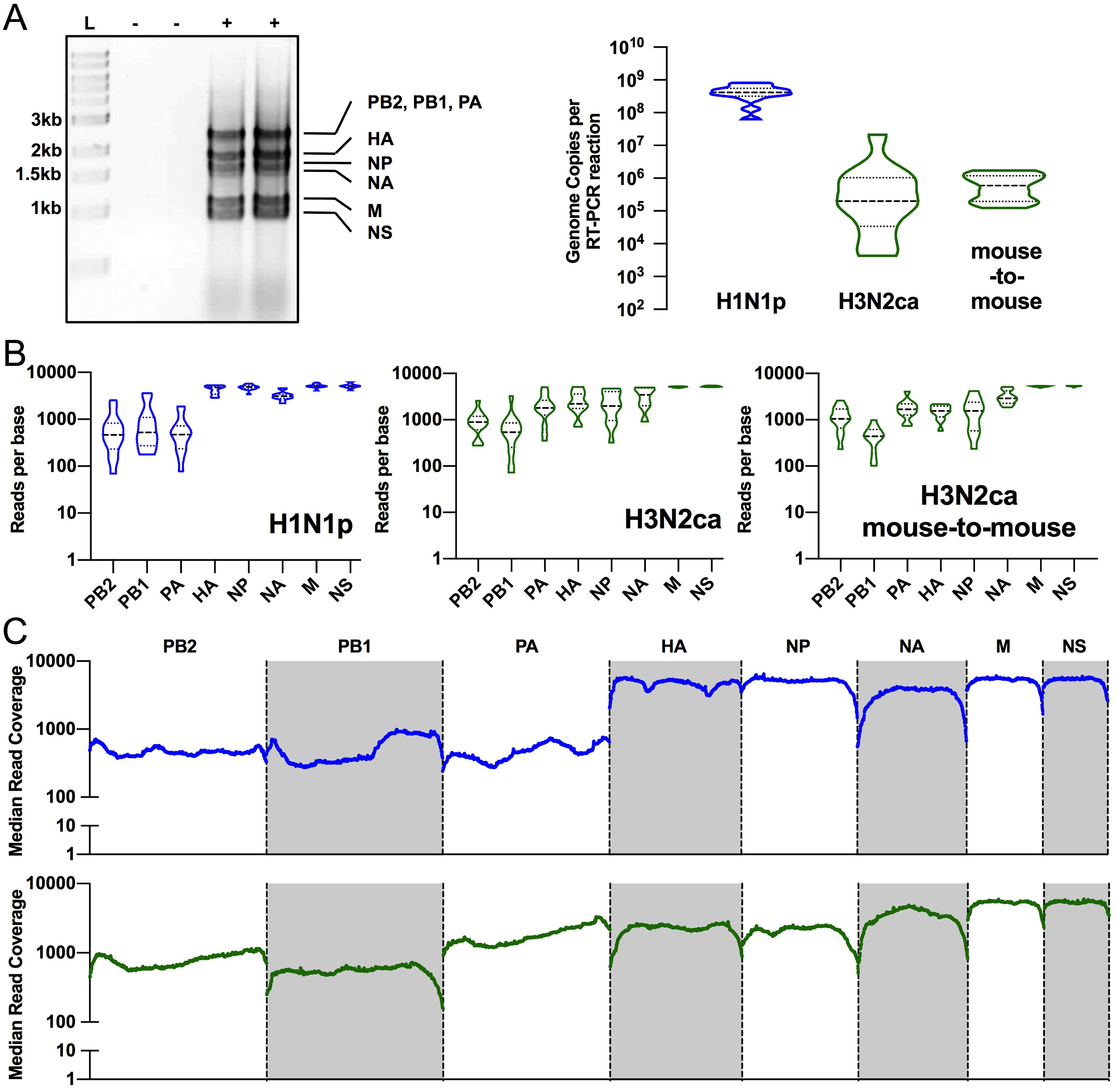
Whole-genome deep sequencing metrics and quality control. (A) Gel migration of products of IAV whole-genome RT-PCR shows specific bands of 8 genome segments. Products were only present in infected lung homogenates (+), absent in uninfected controls (−). RT-PCR reactions generally contained >100,000 input genome copies. (B) Reads across genome segments were not equal, with frequent bias towards smaller segments. (C) Median reads across the genome segments show some variation in coverage across PB2/PB1/PA segments.

### Passage of H1N1p in mice resulted in only a small number of polymorphic sites

Analysis of the H1N1p viruses at each passage in mice showed only very low levels of sequence variation (**Fig. 6**). The plasmid sequence showed no single nucleotide variants (SNVs) above 0.2% at any position, confirming the low errors associated with library preparation and Illumina sequencing. After three passages in the MDCK cells the H1N1p viruses contained just 9 SNVs at >1% of reads in the PB1, PA, HA, and NA segments, and two (in HA) were present at ∼5-7% (**Fig. 6**). We deep sequenced the genomes of the virus in the lungs pooled from each group of mice at each passage in each series, and found no more than 22 unique SNVs (>1%) across the genome in the final passage of each group. Although mutations were present in multiple genome segments, most were found in the PB1 and HA genes. However, even these SNVs represented <5% of the sequences at these positions and none increased significantly during either of the repeated passage series conducted (**Fig. 6**). The two positions that showed a SNV frequency >5% were PB1-Val709 and HA-Asp222 (**Fig. 6**, annotated). The PB1-Val709 locus showed a non-synonymous mutation to Ile (by G2149A) in ∼12% of reads in the first mouse infection of one iteration of the experiment, but then fell to ∼4% by passage 3. The HA-Asp222 locus showed variation in only one passage series, and increased in frequency during each passage. In wild-type mice this locus contained a mixture of two SNVs (nucleotides G747A to code for 222Asn, and A748G to code for 222Gly), while in *CMAH*^−/−^ mice only the Asp222Gly variant was present. Besides these changes, other low-level mutations included both synonymous and non-synonymous changes (**Fig. 6**), but did not include any positions known to effect viral functions.

**Figure 6.**
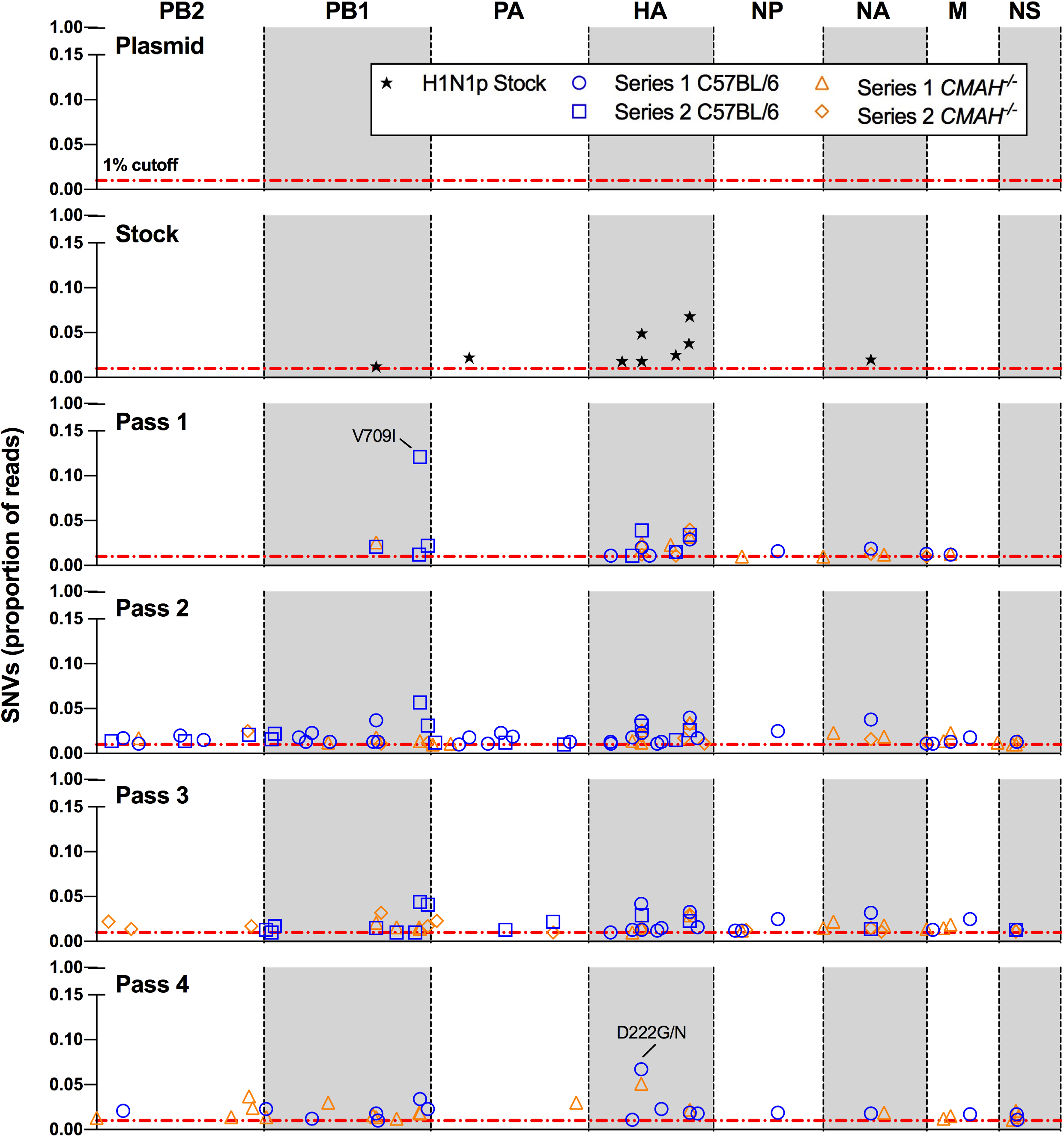
Mutational frequency during H1N1p experimental passage in pooled mice groups. Single nucleotide variants (SNVs) are represented along the genome segments for: the plasmid and virus stocks (black stars), C57BL/6 mice of Series 1 (blue circles), *CMAH*^−/−^ mice of Series 1 (blue squares), C57BL/6 mice of Series 2 (orange triangles), *CMAH*^−/−^ mice of Series 2 (orange diamonds). SNVs present at >20% are annotated for their resulting protein sequence changes.

### The H3N2hu did not establish in mice

Recent human seasonal H3N2 viruses have been reported to replicate poorly in many standard cells in culture and in mice (50, 51), due at least in part to a need for multivalent binding to the Sia receptors to allow infection (52, 53). We recovered the H3N2hu virus on MDCK-SIAT cells which have both higher levels of α2,6-linked Sia and higher densities of Sia overall (54), although growth in wild-type MDCK cells was also successful and the virus produced was comparable in titer and viral genome sequence (data not shown). However, after inoculation into C57BL/6 or *CMAH*^−/−^ mice we detected only low levels of viral RNA by RT-qPCR at three days, and did not see viral antigen in lungs of inoculated mice (**Fig. 4B**). The H3N2hu was clearly propagating weakly in the mice, and did not establish sustained infections after inoculation, and no signs of infection of the second passage mice were detected (data not shown).

### Passage of H3N2ca in mice resulted in more polymorphisms, and rapid appearance of a small number of mutations

The H3N2ca prepared in MDCK cells after plasmid transfection showed one non-synonymous mutation in NA (nucleotide G98A, Ala27Thr) that represented ∼15% of the sequences, along with only 7 other minor variants at >1% of reads (**Fig. 7**). This virus replicated well in mice, and several mutations arose to relatively high levels during both passage series in groups of mice (**Fig. 7**). SNVs were seen across all genome segments during both passage series and in both mice backgrounds. Final numbers of unique SNVs (>1%) in H3N2ca groups ranged from 29 to 41, although diversity consistently collapsed during passage series from as many as 3× the number of unique SNVs early in the series compared to the last pass. In particular, segments PB2, PA, HA, and NA were most likely to have variants that reached over 10% or that approached fixation. The NA-Ala27Thr change present in the inoculum appeared during both passage series in mice. Several other specific mutations arose during the different passage series or mouse genetics cohorts in the PB2, PA, HA, and NA segments (**Fig. 7**, annotated). Three of these mutations rose to high frequency, or near fixation, during passage: PB2-Ser286Gly, PA-Tyr112Cys, and the culture-derived NA-Ala27Thr. Other mutations that arose during passage in mice did not reach 50%, and occasionally fell off during the later passages possibly due to competition with the mutation-containing genomes that reached higher levels. To examine whether the middle-level mutants were arising in single mice among the groups of mice where we had examined pooled samples, we isolated individual lung homogenates from the three P1 mice in Series 2 and passaged each in an additional series of three single mouse-to-mouse lineages (**Figs. 2C, 8**). This revealed that individual mouse passaged viral lineages acquired mutations in the same genome segments (PB2, PA, HA, NA) as were seen in the pooled mouse samples. The NA-Ala27Thr mutation was again rapidly selected in all lineages, but there was no obvious convergent evolution of the other specific mutations. Individual lineages also contained mutants in the NS segments (lineages a and d) that were not observed in group passage series, and some mutants in HA also rose to higher frequency in individual lineages (b, f) than in the group series (**Fig. 8**, annotated).

**Figure 7.**
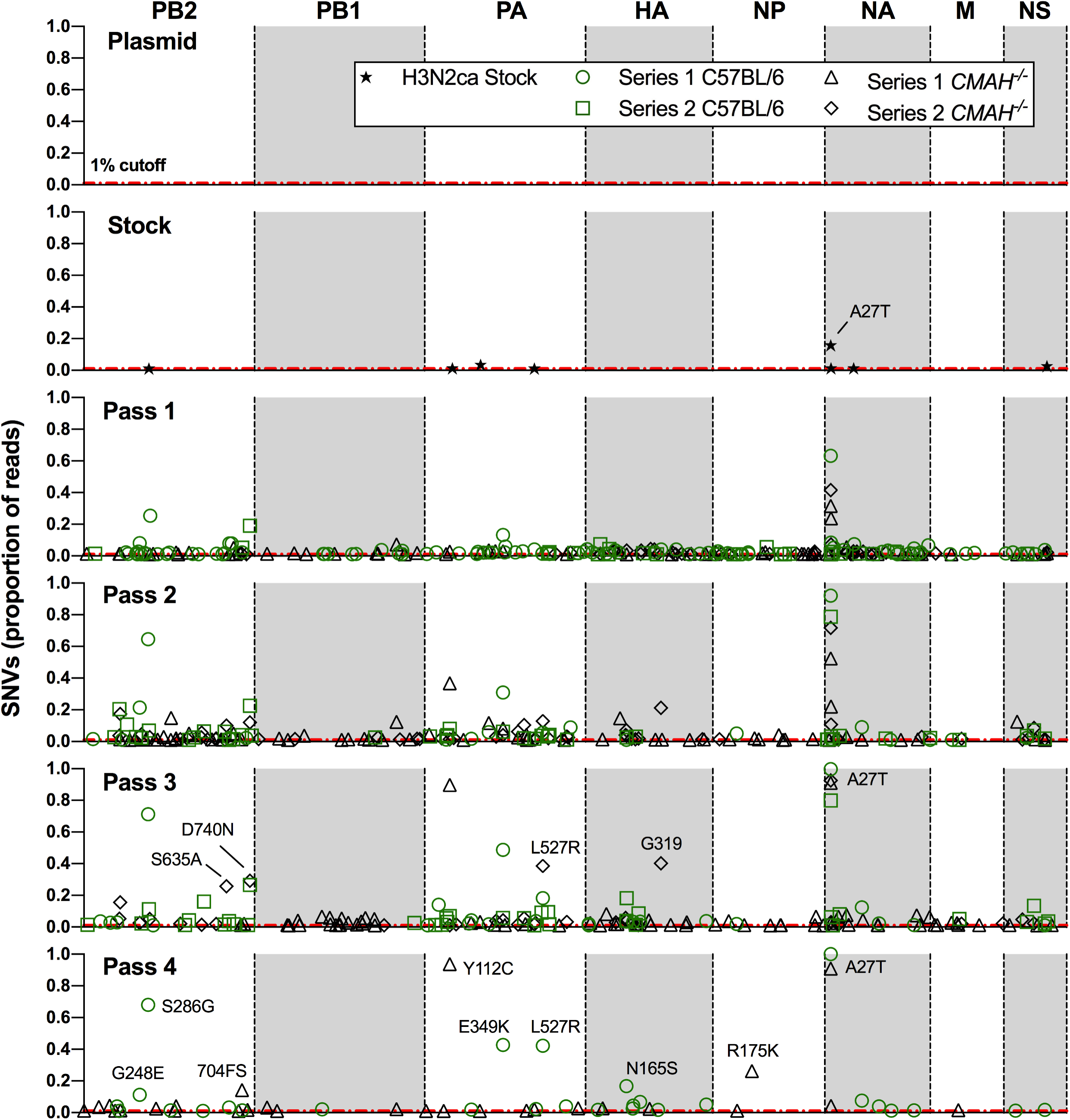
Mutational frequencies during H3N2ca experimental passage in pooled mice groups. Single nucleotide variants (SNVs) are represented along the genome segments for: the plasmid and virus stocks (black stars), C57BL/6 mice of Series 1 (green circles), *CMAH*^−/−^ mice of Series 1 (green squares), C57BL/6 mice of Series 2 (black triangles), *CMAH*^−/−^ mice of Series 2 (black diamonds). SNVs present at >20% are annotated for their resulting protein sequence changes.

**Figure 8.**
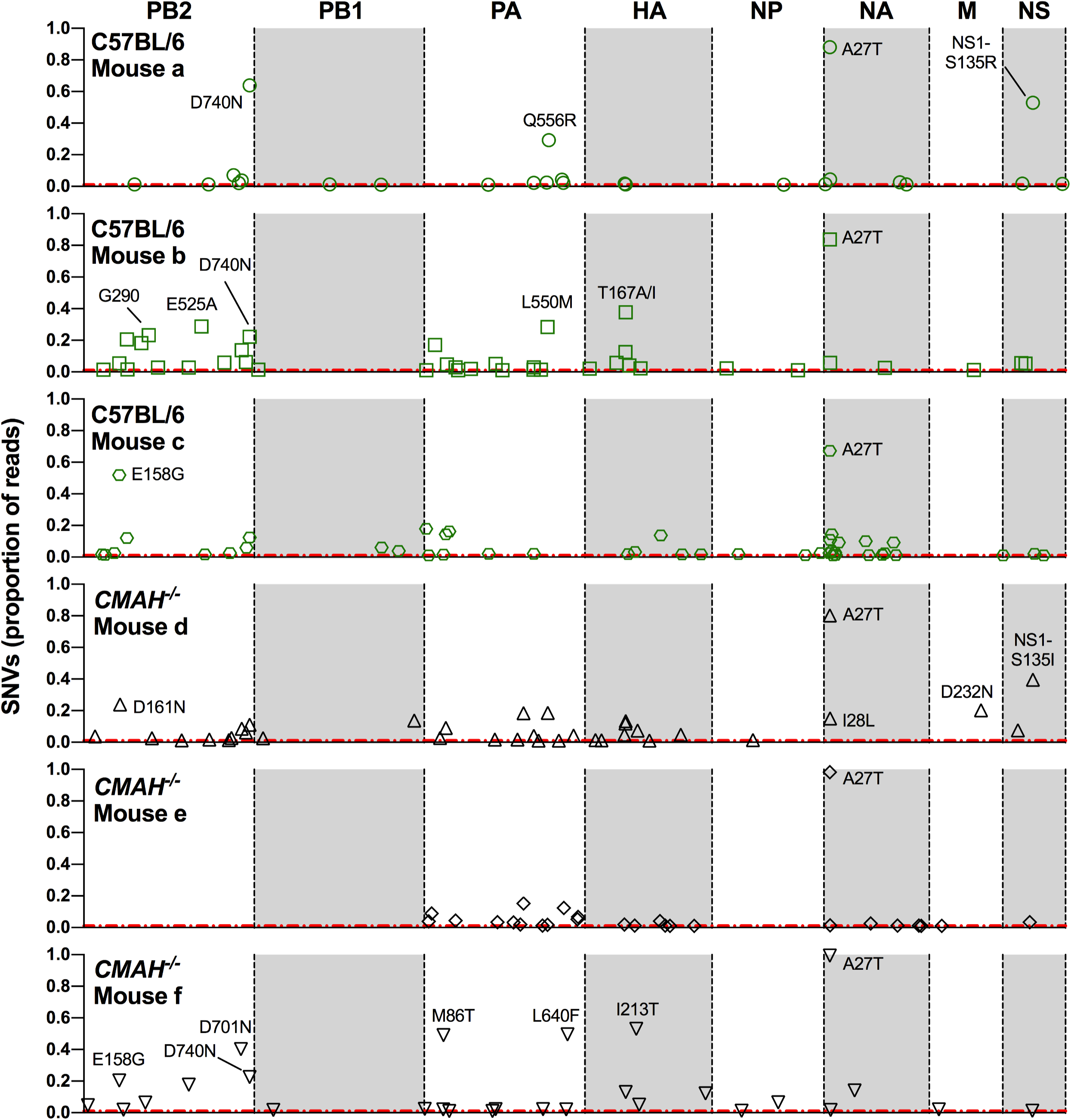
Mutational frequency at the conclusion (passage 3) of H3N2ca mouse-to-mouse experimental passages in C57BL/6 or *CMAH*^−/−^ mice. Single nucleotide variants (SNVs) are represented along the genome segments for: C57BL/6 lineages a (green circles), b (green squares), c (green hexagons), and *CMAH*^−/−^ lineages d (black triangles), e (black diamonds), f (black inverted triangles). SNVs present at >20% are annotated for their resulting protein sequence changes. FS = frame-shift.

## DISCUSSION

We sought to better understand the host-specific variation and evolution of two human- and one canine-adapted IAVs during infection and passage in mice, providing information central to understanding the evolution and potential adaptation of different influenza viruses in this commonly used experimental host. While mouse passaging of many different mammalian and avian IAVs have been reported, this work is novel in the use of complete genome deep sequencing to directly compare the variation of different viruses and for testing the specific role of the modified Sia (Neu5Gc) present in mouse tissues at high levels (but which is absent from humans, ferrets, and most dog breeds). Each virus examined was already adapted to mammals, so the main selection pressures would likely be related to host-specific differences of the mice strain tested, as well as to the experimental lung-to-lung transmission route. To allow us to track newly-emerged variation in the mice, each virus was initiated from reverse genetics plasmids that were also deep sequenced, providing a baseline for comparison. Viral stocks used to initiate the passage series in the mice were prepared by three passages in either MDCK cells (H1N1p and H3N2ca), or in MDCK-SIAT cells which expressed higher levels of α2,6-linked Sia (H3N2hu). To specifically examine the sources of variation in the different virus:mouse passage series, the various passage combinations were designed based on the preliminary data obtained.

### The presence of the non-human modified Neu5Gc Sia did not alter the IAV variation in mice

About 50% of the Sia present in the trachea and lungs of wild-type C57BL/6 mice was Neu5Gc, while *CMAH*^−/−^ mice expressed 100% Neu5Ac, similar to what is seen in humans, ferrets and Western dogs (31, 55–57). The variant Sia measured here are most likely components of both the mucus present in those tissues, as well as on the cell surfaces where they act as receptors for infection. We found no significant effect of the presence of Neu5Gc in mouse respiratory tissue in this study. It is likely that the Neu5Gc Sia does not bind the HA of the viruses tested, so that those do not play a role in IAV infection. In the small number of cases where this has been examined it appears that circulating IAVs preferentially bind the Neu5Ac receptor, and largely ignore the abundant Neu5Gc present in some hosts - as was seen in swine H3N2 strains (58). In Neu5Ac-binding strains where HA is mutated (at position 155) to allow Neu5Gc binding, their NA cleavage is generally inefficient on that Sia, disrupting HA-NA balance and blocking productive infection (59). Our data suggested that the HA binding properties of the viruses tested here are under weak or no selection by Neu5Gc in wild-type mice. The natural ecology of IAV may only involve Neu5Gc binding by viruses infecting hosts that express high levels of that Sia (60–62). Recent work shows that only a small number of avian variants or equine strains demonstrate true Neu5Gc binding by their HA, yet with 2 to 4 fold lower NA specificity for Neu5Gc, including avian N1 and N9, and human N1 (45). Human IAV may evolve to avoid HA binding of inhibitory Sia variants found in other hosts but that appear to be lacking from humans, such as the case of 4-O-acetyl modified Sia present in horse serum (63, 64).

### Deep Sequencing enables a dynamic examination of population variation during evolution

The plasmid sequences showed that the library preparation and Illumina sequencing contributed no variation to the data obtained. The sequence data from the MDCK-passaged virus stocks also showed very few polymorphic positions above 1 or 2% of the total, apart from the NA-Ala27Thr substitution of H3N2ca. The sequences of H3N2ca viruses directly recovered from infected dogs show that Ala is the most common wild-type residue at that position, none show Thr, and a single virus from Korea that had been passaged in culture showing Gly (Genbank AFN06542) (65). This mutation lies within the transmembrane domain of NA and does not have a described function.

### Patterns of evolution of IAVs during mouse passage are strain/subtype contingent

The three viruses showed distinct patterns of evolution when passaged in mice. This is consistent with the ideas around historical contingency, in which related populations will experience distinct evolutionary trajectories despite following the same challenge – in this case mouse host adaptation (66, 67). The failure of H3N2hu (A/Wyoming/03/2003) to replicate appears to be consistent with multiple observations showing that mice are difficult to infect with more recent seasonal H3N2 strains from humans (51). Recent H3N2 strains show significant adaption to human respiratory Sia receptors that are highly branched or multi-antennary N-glycans, likely due to modulated HA avidity in balance with antigenic selection on the epitopes near the top of the HA trimer, close to the Sia binding site (53). In these studies we saw no differences in the sequences of H3N2hu viruses produced in the MDCK-SIAT versus wild-type cells. These results highlight the important role of strain and of subtype sequences on the infection of the same animal, and on the evolution of host adaptation.

Passage of H1N1p repeatedly in either mouse strain revealed only a few positions with low levels of SNV polymorphism, most frequently in the PB1 and HA gene segments. Most polymorphisms were below 5%, while the two mutations present above 5% were each seen in a different iteration of the experiment. The mutations detected included both synonymous and non-synonymous changes. In previous studies of H1N1p various mutations were reported to be present after mouse passage, including in PB2, PA, HA, and NP (68, 69). Only the HA-Asp222Gly/Asn variant corresponded with mutations previously seen in mouse adaptation studies or selected in other hosts. Changes in that position have been seen in human clinical samples and associates with varied pathogenic outcomes in both humans and mice due to changes in receptor specificity by the Sia α2,3 or α2,6 linkages (70, 71). Our observations of little change in mutational frequencies, along with minimal convergence of sequences in repeated studies, suggest that changes seen in mouse passage of H1N1p may be random polymorphisms and hence not directly impact fitness. It is possible that differences between our results and those of previous studies reflects different viral sources, different passaging schemes, or the mice strains used (BALB/c or DBA in some previous studies versus C57BL/6) (72, 73). Interestingly, the deep sequencing of H1N1p passages in C57BL/6 mice in unrelated studies also revealed unique variants and minimal signatures of selective fixation (R. Honce and S. Schultz-Cherry, personal communication).

The speed of adaptation in novel environments is impacted by the fitness effects of individual mutations (74). The H1N1p virus may already be relatively well adapted to mice, so that most mutations do not provide much fitness benefit. Other things being equal, variants with relatively small effect sizes will fix slower even if they are common (75, 76). The mutation rate of IAVs has been estimated at greater than 1 × 10^−4^ substitutions per nucleotide per strand copied (s/n/r), although most of the mutations that arise appear to be removed by purifying selection or lost during bottlenecks that occur during cell-to-cell infection or host-to-host transmission (77). The experimental passages of the virus in our studies did not have tight bottlenecks, which led us to expect robust adaptation if natural selection on mutations of different fitness was present. While many of the variants in the H1N1p viruses did not rise in frequency, the polymorphisms that were observed had arisen *de novo* during the growth of the plasmid-derived viruses in culture and mouse passage, and their maintenance in each virus population was consistent with random sampling effects and perhaps genetic hitchhiking.

H3N2ca inoculation in mice gave rise to considerably more SNV diversity that H1N1p, and the emergence of polymorphisms in PB2, PA, HA, and NA, including near fixation of PB2-Ser286Gly, PA-Tyr112Cys, and NA-Ala27Thr. Some other mutations that rose in frequency, often to >40% of reads in the population, suggest they could be on a path of selective fixation in a new background. Unlike past work in the H3 background (human A/HongKong/1/1968/H3N2), we did not see frequent mutations in HA, nor the PB2-Asp701Asn mutation that was strongly associated with mouse pathogenesis and mammalian adaptation by importin-α interaction and nuclear import of vRNPs (78–80). Other H3N2ca virus mutations that arose in our experiment that parallel past H3 mouse passage observations included PB2-Asp740Asn, PA-Gln556Arg and M-Asp232Asn, but these were mostly seen in single mouse-to-mouse lineages (81). It is possible that the NA-Ala27Thr mutation (of apparent strong fitness advantage but unknown function) influenced the evolutionary path of some of our H3N2ca lineages.

Populations evolving with a larger mutational load may be subject to stochastic outcomes without a strong directed selection coefficient (82). This may explain our results with H3N2ca, in which mutations frequently arose and also increased in frequency, but with little to no convergence of specific mutations between several repeat iterations of the same experiment (apart from the NA-Ala27Thr culture-selected mutation already present in all passage series). Given sufficiently large population sizes and turnover of the virus, evolving populations will rapidly fix highly beneficial mutations, which could converge if related to a strong and focused selective pressure (83); that this does not occur here again suggests that there is likely no central selective force/constraint of mouse host adaptation.

Overall, this study shows that IAV host adaptation in mice is highly contingent on the specific virus used in the experiment, may exhibit only weak signatures of natural selection with little convergent enrichment of polymorphisms towards population fixation. Greater comparison of mouse passage experiments between different laboratories, along with analysis of key population dynamic variables and deep sequencing, will be useful in explaining the host-specific growth of different viruses in mice.

## MATERIALS AND METHODS

### Cells and viruses

MDCK cells were obtained from the American type culture collection (ATCC, CCL-34). Variants of those MDCK cells with increased levels of α2,6-linked Sia cells were prepared in the laboratory by transfection with the ST6Gal1 gene in a plasmid under the control of the CMV promoter (pcDNA3.1, Invitrogen). Cell clones with increased levels of α2,6-linked Sia were identified by staining with the *Sambucus nigra* (SNA) lectin, and termed MDCK-SIAT cells. HEK293T cells were obtained from ATCC (CRL-3216). All cells were grown in DMEM with 10% fetal calf serum, and 50µg/ml gentamycin.

Three IAV strains were derived from reverse genetics plasmids, comprising (i) human H1N1 pandemic IAV (A/California/04/2009, H1N1p) in plasmid pDP2002, (ii) human H3N2 seasonal IAV (A/Wyoming/3/2003, H3N2hu) in plasmid pDZ, and (iii) a canine H3N2 IAV (A/Canine/IL/11613/2015, H3N2ca) in pDZ. The plasmid encoding each viral segment was prepared from a single bacterial colony, and an 8-plasmid mixture for each virus was prepared and used for transfection of a 3:1 co-culture of HEK293T cells and MDCK cells, while for HEK293T and MDCK-SIAT cells for H3N2hu were found to be required to prepare and propagate the virus. Each virus was passaged two additional times in the same MDCK, or MDCK-SIAT variant cells, to generate a passage-3 stock, which was titered for infectivity by TCID_50_ assay, and for viral RNA titer by quantitative reverse transcriptase PCR (RT-qPCR). Each plasmid mixture (as DNA) and the resulting virus stocks (from RNA) were then used to generate libraries for Illumina sequencing, as described below, revealing the original sequences and any baseline variation of the viruses used to start the mouse inoculations.

### Mouse inoculation and serial passage

All mouse studies followed protocols approved by the Cornell University Institutional Animal Care and Use Committee. The passage series and analysis of each virus is diagrammed in **Fig. 2**. Control wild-type (C57BL/6) and *CMAH*^−/−^ (B6.129X1-*Cmah^tm1Avrk^*/J) mice were obtained from Jackson Laboratories and housed and/or bred on site. Mice (aged 6-10 weeks) were anesthetized using isoflurane gas and inoculated intra-nasally with 50µl of 10^4^ TCID50 units (MDCK or MDCK-SIAT) of each virus in PBS. Mice were observed and weighed each day, euthanized 3 days post-infection (dpi) and tissues harvested. Single lung lobes of each mouse averaged ∼100mg, and were homogenized with 0.5ml of added sterile PBS (∼20% w/v). Homogenates were clarified by centrifugation at 1,000 × *g* for 10 min. Individual mice sample lung homogenates were stored at −80°C. In some studies lung homogenate supernatants (n =4 or 3 of like virus strain and mouse genetics) were also pooled and aliquoted, and 50µl used to inoculate each mouse in the next passage.

As the H3N2ca-inoculated mouse samples showed the greatest number and level of mutations, and to understand the dynamics and variation within individual mice, for the second passage series lungs of individual mice were passed on to additional individual mice to reveal any differences in the dynamics of single mouse-to-mouse lineages (**Fig. 2C**).

### Viral RNA extraction and virus copy number quantitation by qPCR

Viral RNA (vRNA) was isolated from virus-infected lung homogenate supernatant using the QIAamp Viral RNA Mini Kit (QIAGEN). Influenza genome copies were quantified from RNA isolations by reverse transcription and quantitative PCR (RT-qPCR) for the M segment modified from CDC protocol (84). Products were amplified using Path-ID (Applied Biosystems) with M-specific primers (5’ to 3’, F: GACCRATCCTGTCACCTCTGAC, R: AGGGCATTYTGGACAAAKCGTCTA), probed with 5’-TGCAGTCCTCGCTCACTGGGCACG-3’ and run on a 7500 Fast Real-Time platform against a standard curve.

### Library generation and NGS sequencing

Influenza whole-genome reverse transcription and polymerase chain reaction (RT-PCR) amplification of cDNA from all 8 viral genome segments was performed using a modification of Zhou et al. (85). Total vRNA was incubated with reaction buffer, Superscript III and Platinum Taq-HiFi (Invitrogen) in the presence of universal influenza amplifying primers (5’ to 3’, uni12a: GTTACGCGCCAGCAAAAGCAGG; uni12b: GTTACGCGCCAGCGAAAGCAGG; uni13: GTTACGCGCCAGTAGAAACAAGG). RT-PCR reactions were performed as first-strand synthesis (42°C) for 50 min, followed by 5 cycles at 42°C annealing, then by 33 cycles at 57°C annealing. Viral cDNA products were purified by Agencourt AMPureXP magnetic beads (Beckman Coulter), eluted in Tris buffer and quantified by QuBit (Invitrogen). Amounts of 1 ng cDNA were used to prepare NGS sequencing libraries using Nextera XT DNA Library Prep Kit (Illumina), with unique index adaptors for each sample. Pooled libraries were run on an Illumina MiSeq v2 for 250bp paired reads in the Cornell Animal Health and Diagnostic Center Molecular Diagnostics Lab or the Cornell Genomics Facility of the Biotechnology Research Center.

### Sequence analysis and variant calling

Analysis was performed in Geneious v.11.1.5. Read trimming was performed by the BBDuk script, followed by read merging, and alignment to the reference genomic plasmid sequence for each virus. For variant calling, we considered those at >1% frequency with minimum 500 read coverage (although the sequences appeared generally accurate to 0.2% when at high coverage). Recognizing the potential for PCR-derived errors being observed as SNVs, and our use of single library runs per sample, we focused our analysis on those mutations that maintain passage-to-passage. Longitudinal sampling of SNVs would allow unique samples to act as proxy duplicates and further suggest biological relevance of those mutations.

### HPLC analysis of mouse respiratory tissue variant sialic acids

The Sia composition of the mouse trachea and lung samples was determined by incubating with 2M acetic acid at 80°C for 3 hr, filtration through a Microcon 10 kD centrifugal filter (Millipore), and drying in a SpeedVac vacuum concentrator. Released Sia were labeled with 1,2-diamino-4, 5-methylenedioxybenzene (DMB, Sigma Aldrich) for 2.5 hr at 50°C (86). HPLC analysis was performed using a Dionex UltiMate 3000 system with an Acclaim C18 column (ThermoFisher) under isocratic elution in 7% methanol, 7% acetonitrile, and 86% water. Sia standards included bovine sub-maxillary mucin and commercial standards for Neu5Ac and Neu5Gc (Sigma Aldrich). Statistical analyses were performed in PRISM software (GraphPad, version 8).

### Histochemistry of mouse respiratory tissue for sialic acid distribution and experimental lungs for viral antigen

Expression of α2,3 and α2,6 linked Sia in the trachea and lungs of mice was examined by preparing frozen sections of OCT-embedded tissue. Sections were fixed for 30 min with 10% buffered formalin, then incubated with *Maackia amurensis* (MAA-II, MAH) or *Sambucus nigra* (SNA) lectins conjugated with biotin (Vector Laboratories, Burlingame, CA). Sections were then probed by streptavidin-HRP followed by incubation with the substrate NovaRed (Vector).

The additional lung lobe of each experimental mouse was stored in 10% buffered formalin. Lungs were paraffin-embedded and cut for both H&E staining and immunohistochemistry. The presence of influenza viral antigen in lung tissue was determined by staining with the anti-NP monoclonal antibody (ATCC HB-65, clone H16-L10-4R5), followed by anti-mouse goat IgG conjugated with HRP, and then incubation with the substrate NovaRed.

### Graphing and analysis

All figure graphs were generated in GraphPad Prism v.8.1.2.

## ACKNOWLEDGEMENTS

Reverse genetics plasmids for A/Wyoming/3/2003/H3N2 were provided by Adolfo García-Sastre at the Icahn School of Medicine, Mount Sinai. Reverse genetics plasmids for A/California/04/2009/H1N1 were provided by Daniel Perez at the University of Georgia. We thank support services at the Cornell College of Veterinary Medicine Animal Health Diagnostic Center (AHDC): the Molecular Diagnostics Lab for support in RT-qPCR analysis as well as Illumina-MiSeq protocol development and sequencing runs, and the Histology lab for support in FFPE tissue-embedding and slide preparation. This work was supported in part by CRIP (Center of Research in Influenza Pathogenesis), an NIAID funded Center of Excellence in Influenza Research and Surveillance (CEIRS) contract HHSN272201400008C to CRP, and by NIH grant R01 GM080533 to CRP. IEHV was supported by NSF award DGE-1650441. ECH is supported by an ARC Australian Laureate Fellowship (FL170100022).

